# State-of-the-art image and video quality assessment with a metric based on an intrinsically nonlinear neural summation model

**DOI:** 10.1101/2022.12.22.521587

**Authors:** Raúl Luna, Itziar Zabaleta, Marcelo Bertalmío

## Abstract

The development of automatic methods for image and video quality assessment that correlate well with the perception of human observers is a very challenging open problem in vision science, with numerous practical applications in disciplines such as image processing and computer vision, as well as in the media industry. In the past two decades, the goal of image quality research has been to improve upon classical metrics by developing models that emulate some aspects of the visual system, and while the progress has been considerable, state-of-the-art quality assessment methods still share a number of shortcomings, like their performance dropping considerably when they are tested on a database that is quite different from the one used to train them, or their significant limitations in predicting observer scores for high framerate videos. In this work we propose a novel objective method for image and video quality assessment that is based on the recently introduced Intrinsically Non-linear Receptive Field (INRF) formulation, a neural summation model that has been shown to be better at predicting neural activity and visual perception phenomena than the classical linear receptive field. Here we start by optimizing, on a classic image quality database, the four parameters of a very simple INRF-based metric, and proceed to test this metric on three other databases, showing that its performance equals or surpasses that of the state-of-the-art methods, some of them having millions of parameters. Next, we extend to the temporal domain this INRF image quality metric, and test it on several popular video quality datasets; again, the results of our proposed INRF-based video quality metric are shown to be very competitive.

## 1. INTRODUCTION

Image quality evaluation is of crucial importance in the media industry, where it has numerous practical applications, but also in applied disciplines such as image processing and computer vision, where it plays an important role in the development, optimization, and testing of algorithms. It may also be used to optimize any trade-offs between the components of an image/video transport system (i.e. video compression ratios, reserved network bandwidth) and perceived quality, aiming for a good user experience while maximizing the efficiency of the transmission, and thus reducing the ecological footprint associated to streaming. Subjective evaluation, consisting in measuring image quality by human beings, is costly and time-consuming. Therefore, the goal of objective quality assessment is to develop automatic methods that produce quantitative measures that are consistent with the perception of human observers. But this is a very challenging open problem in vision science, given the limitations of current visual perception models and the way they are exacerbated by emerging image display technologies of ever-increasing resolution, contrast, color gamut and framerate (Bertalmío, 2019).

Image quality methods can be divided into three categories: full-reference methods, which compare an original image with a distorted version of it; reduced-reference methods, that compare some characteristics of the distorted and reference image since the complete reference image is not available; and no-reference methods (also called blind models), operating solely on the distorted image.

In this article we will focus on full-reference methods, which constitute the vast majority of image quality approaches. A simple solution, and possibly the most widely used metric to estimate image quality, is the peak signal-to-noise ratio (PSNR), which is a nonlinear transform of the mean square error (MSE) between the reference and the distorted images, another very popular metric. These metrics are simple to calculate, and they have a clear physical meaning; however, they are not very well correlated with perceived visual quality (Wang et al., 2004).

Therefore, in the last two decades, the goal of image quality assessment (IQA) research has been to improve these metrics by developing more sophisticated methods that mimic some aspects of the visual system. For instance, the Normalized Laplacian Pyramid Distance (NLPD) (Laparra et al., 2016) is based on transformations present in the early visual system: local luminance subtraction and local gain control, obtained from a decomposition of images using a Laplacian pyramid; the Structural Similarity Index (SSIM) (Wang et al., 2004) is based on the hypothesis that the human visual system is highly adapted for extracting structural information from the viewing field; the Feature Similarity Index (FSIM) (Zhang et al., 2011) is based on the assumption that the human visual system understands an image according to its low-level features, such as the phase congruency, which measures the significance of a local structure, and the image gradient magnitude, which encodes contrast information; the Visual Signal-to-Noise Ratio (VSNR) (Chandler & Hemami, 2007) analyses visual perception distortions in the wavelet domain; the Noise Quality Measure (NQM) (Damera-Venkata, Kite, Geisler, Evans & Bovik, 2000) is based on the contrast pyramid by Peli (1990); and the Visual Information Fidelity Measure (VIF) (Sheik & Bovik, 2006) is based on natural scene statistics and models of the image degradation process and the human visual system. There are also learning-based methods that learn a metric from a set of training images and their corresponding perceptual scores. For instance, the Learned Perceptual Image Patch Similarity (LPIPS) metric (Zhang et al., 2018) is based on the hypothesis that perceptual similarity is a consequence of visual representations, as the authors found that internal activations of networks trained on high-level image classification tasks correspond well to human perceptual judgments; another example is the Deep Image Structure and Texture Similarity (DISTS) Metric (Ding et al., 2020), which uses a variant of the VGG convolutional neural network to construct a function that combines structure and texture similarity measurements between corresponding feature maps of the reference and distorted images; and PerceptNet (Hepburn et al., 2020), which is a convolutional neural network where the architecture reflects the structure and various stages in the human visual system: a cascade of canonical linear filters and divisive normalization layers simulate the retina-LGN-cortex pathway.

For video quality assessment (VQA), a simple option is to apply image quality metrics on a frame-by-frame basis, but this type of approach often provides a limited performance, especially in the case of high framerate (HFR) videos (Madhusudana et al., 2021). Therefore, the state-of-the-art in VQA are algorithms specifically developed for video, including the Spatio-temporal Reduced Reference Entropic Differences (ST-RRED) (Soundararajan & Bovik, 2012) or the Spatial Efficient Entropic Differencing for Quality Assessment (SpEED) (Bampis et al., 2017), both measuring quality deviations by computing spatial and temporal entropic differences in the band-pass domain; Frame Rate dependent Quality Metric (FRQM) (Zhang et al., 2017), which outputs quality measurements by calculating absolute differences between sequences that have been temporally filtered by a wavelet; Video Multi-method Assessment Fusion (VMAF) (Li et al., 2016), which, using a Support Vector Regressor, fuses a frame-difference feature with a detail feature and with features obtained from a Visual Information Fidelity (VIF) measure (Sheikh & Bovik, 2006); Deep Video Quality Assessor (deepVQA) (Kim et al., 2018), which combines a CNN model with a Convolutional Neural Aggregation Network (CNAN) used for temporal pooling; the Visual Quality Metric (VQM) (Pinson & Wolf, 2004), which uses reduced-reference technology (ITU-T, 2005) to provide estimates of video quality; the perceptual spatio-temporal frequency-domain based MOtion-based Video Integrity Evaluation (MOVIE) index (Seshadrinathan & Bovik, 2009), which monitors distortions that are perceptually relevant along motion trajectories; or the more recent Generalized Spatio-Temporal Index (GSTI) (Madhusudana et al., 2020), which calculates entropic differences between responses that have been temporally band-pass filtered.

Despite the notable advances in the field, it is important to point out that state-of-the-art quality assessment methods still share a number of shortcomings, like their performance dropping considerably when they are tested on a database that is quite different from the one used to train them (Ding et al., 2021), or their significant difficulties in predicting observer scores for HFR videos (Madhusudana et al., 2021). In this study we aim to overcome these limitations by estimating perceived image and video quality using a model for neural summation introduced recently, called the Intrinsically Non-linear Receptive Field (INRF) (Bertalmío et al., 2020) formulation; the INRF model successfully explains experimental data that linear receptive field models are unable to explain or do not explain accurately (Bertalmío et al., 2020), and it has been shown to be very promising as a tool to develop IQA methods given its ability to model complicated perceptual phenomena.

The main contributions of this work are as follows. Firstly, we start by optimizing, on a classic image quality database, the four parameters of a very simple INRF-based metric, and proceed to test this metric on three other databases, showing that its performance equals or surpasses that of the state-of-the-art IQA methods, some of them having millions of parameters. Secondly, we extend to the temporal domain this INRF image quality metric, and test it on several popular video quality databases; our results show that the proposed INRF-based VQA is very competitive, ranking best in several challenging scenarios like those provided by a very recent dataset for high frame rate videos. Finally, and to the best of our knowledge, the approach of using a neural summation model to create IQA and VQA methods is completely novel, and given its success it might pave the way for other neuroscience models to inform the design of new image quality assessment algorithms.

The structure of this manuscript is as follows. Section 2 explains the INRF model and how it is used for IQA and VQA. Section 3 provides details on how the INRF parameters are optimized for IQA and VQA usage, describes the different datasets on which performance is tested and explains further points on how the assessment of INRF for IQA and VQA models is performed. Section 4 shows the performance of INRF, employed for IQA and VQA, on different datasets, and results are compared with performance of other state-of-the-art IQA and VQA algorithms on those same datasets. Finally, in Section 5, results and their implications are discussed.

## 2. PROPOSED METHODS FOR IQA AND VQA

### 2.1. Overview of the INRF neural summation model

In vision science, the receptive field (RF) of a neuron is the extent of the visual field where light influences the neuron’s response. In the “standard model” of vision, the first stage is a filtering operation consisting of multiplying the intensities at each local region of an image stimulus by the values of a filter (the weights of the RF), and summing the resulting intensities (Carandini et al., 2005); this weighted sum may then be normalized by the responses of neighboring neurons and passed through a point-wise nonlinearity. Many scientists have come to accept this linear-plus-nonlinear (L+NL) formulation as a working model of the visual system (Olshausen & Field, 2005), both in visual neuroscience and in visual perception, and while there have been considerable improvements on, and extensions to, the standard model, the linear RF remains as the foundation of most vision models. But there are a number of problems that are inherent to considering the RF as having a linear form, of which we will highlight three:

- adaptation makes the linear RF change with the input and, in fact, the linear RF has been observed to have different sizes, orientations, preferred directions or even different polarity (ON/OFF) for different stimuli (Cavanaugh et al., 2002; Coen-Cagli et al., 2012; Jansen et al., 2018);
- a linear RF depends on the choice of basis functions used to estimate it (Vilankar & Field, 2017);
- the linear RF is not supported by more recent neuroscience, and a growing number of neuroscience studies show that in general individual neurons *cannot* be modeled as a linear RF followed by an output nonlinearity (Poirazi et al., 2003; Polsky et al., 2004; London and Häuser, 2005; Silver, 2010; Rodrigues et al., 2021).

In contrast, the INRF formulation is a physiologically-plausible single-neuron summation model which, unlike the linear RF:

- embodies the efficient representation principle and can remain constant in situations where the linear RF must change with the input;
- is a generalization of the linear RF that is much more powerful in representing nonlinear functions;
- is consistent with more recent studies on dendritic computations.

The INRF equation for the response of a single neuron at location *x* (a set of 2D spatial coordinates, horizontal and vertical) is:

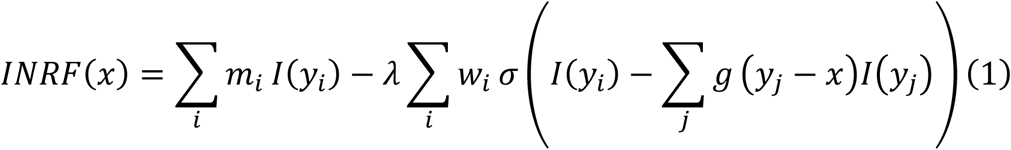

where *m*_*i*_ stands for a 2D kernel *m*(*x, y*_*i*_), locations *y*_*i*_ (vertical and horizontal 2D spatial coordinates) are neighbors of *x, λ* is a scalar, *w*_*i*_ stands for a 2D kernel *w*(*x, y*_*i*_), *σ* represents a non-linearity, and *g* is also a 2D kernel.

The model is based on knowledge about dendritic activity: some dendritic branches act as non-linear units (Silver, 2010), a single non-linearity is not enough to model dendritic computations (Poirazi, Brannon & Mel, 2003), and there is feedback from the neuron soma to the dendrites (London & Häusser, 2005). In the INRF model, some dendrites are linear and their contributions are summed with weights *m*_*i*_, and some other dendrites are nonlinear and their contributions are summed with weights *w*_*i*_. The feedback from the soma is reflected in the shifting nature of the nonlinearity *σ*, expressed by the term Σ_*j*_*g*(*y*_*j*_ − *x*)*I*(*y*_*j*_), which has the effect of making the nonlinearity change for each contributing location *y*_*i*_.

Using a single, fixed INRF module where the kernels *m, w, g* have Gaussian form and the nonlinearity is a power-law sigmoid, and applying it to grayscale images, where *x* and *y*_*i*_ now denote pixel locations and *I*(·) are pixel values, the model response *INRF*(·) emulates the perceived image and can explain several visual perception phenomena that challenge the standard model (Bertalmío et al., 2020):

- The “crispening” effect. Brightness perception curves show “crispening” (slope increase) at the surround luminance level when the background is uniform, but the phenomenon disappears for salt and pepper background (Kane & Bertalmío, 2019). The INRF response qualitatively replicates both cases with a fixed set of parameters, which is not possible with a linear RF formulation.
- White’s illusion under noise. The INRF output qualitatively predicts the observers’ response to White’s illusion when bandpass noise is added, while in Betz et al. (2015) none of the vision models that were tried, based on linear RFs, were able to replicate this behavior.
- Light/dark asymmetry (the *“irradiation illusion”*). This phenomenon can be reproduced with a fixed INRF formulation, while a L+NL model needs to change with the stimulus (Kremkow et al., 2014).

In short, we could say that the INRF formulation, a non-linear transform of light intensity values, is a good estimator of brightness perception. This is a very valuable property when creating an image quality assessment method, as we discuss next.

### 2.2 IQA with the INRF model

Given that applying the INRF model to a grayscale image produces a result that appears to closely resemble how the image is perceived, Bertalmío et al. (2020) proposed a very simple image quality metric: given an image *I*, and its distorted version *I*_*D*_, the INRF transformation is applied to both of the images, obtaining *O* and *O*_*D*_, and then the root mean square error (RMSE) between the processed images is computed. The underlying idea here was that the INRF model performs a sort of “perceptual linearization”, a nonlinear transform that brings *light-intensity* images (whose direct comparison, with metrics like RMSE or PSNR, does not correlate well with perception), to the space corresponding to *perceptual* images (which can now be compared with a simple Euclidean metric like RMSE). This metric had only five parameters that were optimized for the “crispening” brightness perception experiment mentioned above (with *σ* a power-law sigmoid and *g* a delta function) on a handful of synthetic images, and despite this fact and the simplicity of the metric, when tested on the natural image database TID2013 (Ponomarenko et al., 2015) it was shown to have a performance very similar to that of the state-of-the-art deep learning perceptual metric LPIPS (Zhang et al., 2018), with over 24 million parameters and close to 500K human judgements for labeling pair-comparison preferences on the 160,000+ natural images it used for training.

Based on this very promising result, here we propose a full-reference INRF-based IQA method in the following way (see Figure 1A for a graphical explanation of the process):

**Figure 1.**
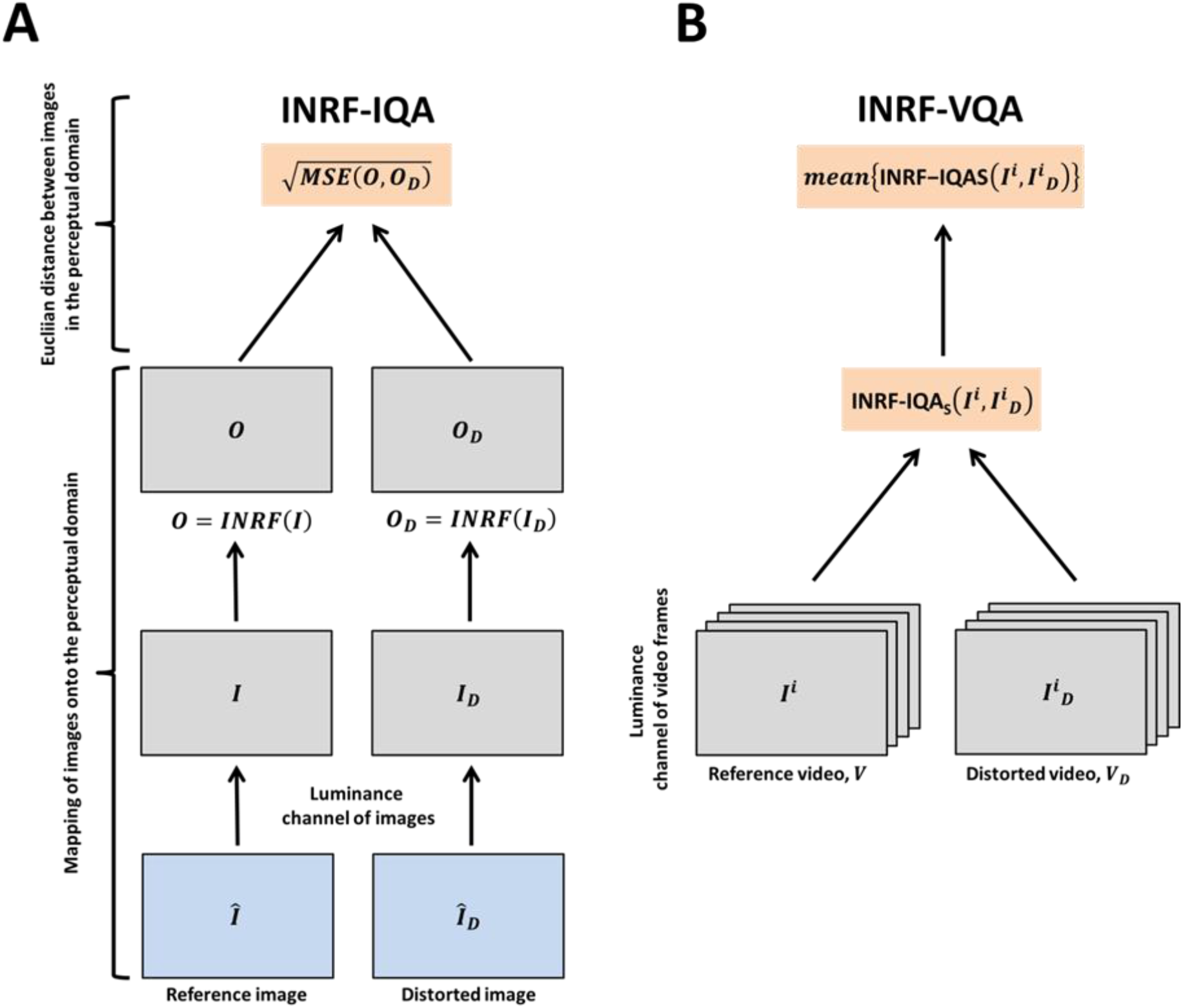
Schematic representation of how INRF-IQA and INRF-VQA are calculated. A. Calculation of INRF-IQA. The luminance component of a reference image, *Î*, and a distorted image, *Î*_*D*_, are mapped onto the perceptual domain by applying an INRF transformation on them: *O* and *O*_*D*_ are respectively obtained. Next, the Euclidean distance of the images in the perceptual domain is calculated by computing the root-mean-squared error of the INRF-transformed images. B. Calculation of INRF-VQA. The process outlined in A is applied on a frame-wise basis to the luminance component of the video frames in a reference video, *V*, and a distorted video, *V*_*D*_. These video frames are referred to as *I*^i^ and *I*^i^_*D*_ respectively. INRF-IQA is calculated using the reference video frames, *I*^i^, and the distorted video frames, *I*^i^_*D*_, that compose the reference and distorted videos, *V* and *V*_*D*_. The final INRF-VQA metric is obtained by computing the mean of the frame-wise INRF-IQA scores.

1. Given a grayscale image *I*, the INRF transformation applied to it produces an image *O* whose value at each pixel location *x* is computed as:

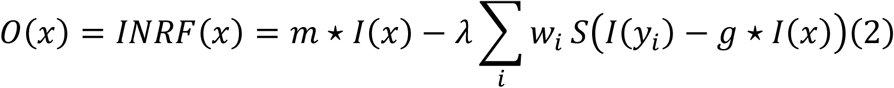

where the kernels *m, w, g* are 2D Gaussians of standard deviations *σ*_*m*_, *σ*_*w*_, *σ*_*g*_ respectively, *λ* is a scalar, *S* is a sigmoid that has the form of an **atan** function, and the symbol *\** denotes the 2D convolution.
2. The INRF-IQA value comparing image *I* and its distorted version *I*_*D*_ is computed as:

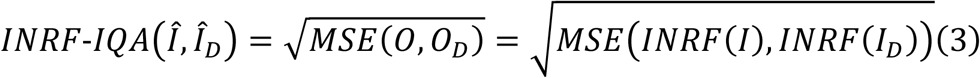

where grayscale image *I* is the luminance channel of *Î*, grayscale image *I*_*D*_ is the luminance channel of *Î*_*D*_, and the INRF transform of an image is computed as in Eq. 2.
3. The values for the four parameters of the metric, namely *σ*_*m*_, *σ*_*w*_, *σ*_*g*_ and *λ*, are chosen so as to maximize the correlation between the INRF-IQA perceptual distance of Eq. 3 and the mean opinion scores (MOS) of human observers over the large-scale natural image database TID2008 (Ponomarenko et al., 2009) (see Methods).

We must note that the IQA method in Bertalmío et al. (2020) used a different nonlinearity (a power-law sigmoid instead of the atan function), considered *g* to be a delta function instead of a 2D kernel, and used as parameter values the same set that allowed the INRF model to reproduce the results of certain brightness perception experiment.

### 2.3. VQA with the INRF model

We propose as a full-reference INRF-based VQA metric a straightforward combination of the INRF-IQA metric of Eq. 3 with a simple temporal pooling strategy: the image metric is applied frame-by-frame, and then the results are averaged (see Figure 1B for a graphical explanation of the process). That is, given a reference video *V* and a distorted version *V*_*D*_, the output of our proposed INRF-VQA metric is:

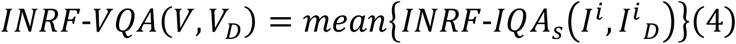

where *I*^i^ and *I*^i^_*D*_ are, respectively, the luminance channel of the i-th frame of *V* and *V*_*D*_, and the IQA metric *INRF-IQA*_*s*_ has the same form as in Eq. 3 and the same value of 3 for the parameter *λ*, but the spatial kernel sizes *σ*_*m*_, *σ*_*w*_, *σ*_*g*_ are scaled by a factor *f* that denotes the ratio between the size of the frames in the video *V* and the size of the images in TID2008 (that were used to optimize *σ*_*m*_, *σ*_*w*_, *σ*_*g*_). For instance, if *V* is a 2K video of resolution 1920*x*1080, and given that images in TID2008 are of size 512*x*384, the scaling factor is 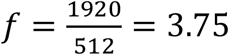.

## 3 METHODS

### 3.1. Optimization details

The parameter values of INRF-IQA (Eq. 3) are found through optimization on the TID2008 dataset (Ponomarenko et al., 2009) to maximize the Pearson correlation coefficient (PLCC) between the observers’ scores for the images in TID2008 and the corresponding INRF-IQA scores. In particular, a grid search optimization method is used to optimize the INRF-IQA parameter values *σ*_*m*_, *σ*_*w*_, *σ*_*g*_ and *λ*, and the optimal parameters found are: *σ*_*m*_ = 1.74, *σ*_*w*_ = 25, *σ*_*g*_ = 1 and *λ* = 3.

### 3.2. IQA datasets

We use the LIVE (Sheikh et al., 2006), CSIQ and TID2013 (Ponomarenko et al., 2015) datasets to test the performance of our INRF-IQA metric on images; each dataset also containing subjective image quality scores from observers for those same images. The LIVE dataset consists of 779 images. It has a total of 29 reference images whose distorted versions are achieved by applying 5 different types of distortion (Gaussian blur, additive white Gaussian noise, JPEG compression, JP2K compression and a simulated fast fading Rayleigh channel) with different distortion levels. The CSIQ dataset has 866 images, with 30 being reference ones. Their distorted versions are obtained through Gaussian blur, Gaussian pink noise, Gaussian white noise, JPEG compression, JP2K compression and contrast change. The TID2013 dataset extends the previous TID2008 one (Ponomarenko et al., 2009). It is composed of 3000 images, out of which 25 are reference ones. These are distorted to achieve the rest of the images by applying 24 distortion types (7 new types of distortions with respect to TID2008) each with 5 distortion levels. Distortion types are rich, spanning from Gaussian noise, Gaussian blur, lossy compression of noisy images, and distortions such as JPEG to more uncommon distortions like non-eccentricity pattern noise.

### 3.3. VQA datasets

We test the performance of our INRF-VQA model on four popular video quality databases, all publicly available and containing observers’ scores for a number of common spatial and temporal distortions: LIVE-YT-HFR, LIVE-MOBILE, LIVE-VQA and VQEG-HD3.

The very recent LIVE-YT-HFR dataset (Madhusudana et al., 2021) spans 16 different video categories, each showing a different progressively scanned natural scene. Out of those 16 contents, 11 have 2K spatial resolution and 5 have 4K resolution. Within each of the video contents, videos with different frame rates exist, out of which we use those of 120 fps, 60 fps and 30 fps. Each video content with a given frame rate has 5 possible compression levels (FFmpeg VP9 compression [Mukherjee et al., 2015], and single-pass encoding varying the Constant Rate Factor [CRF]). For instance, a given content and its videos with a given frame rate sum a total of 5 videos: 1 video with lossless compression (CRF=0) plus 4 videos with compression levels ranging from CRF=4 to CRF=63. Finally, for each video content, the 120 fps video with lossless compression (CRF=0) is referred to as the reference sequence. All the remaining videos within a content (that is, 120 fps videos with CRF values larger than 0, and 60 fps and 30 fps videos), are the distorted sequences.

The LIVE-MOBILE dataset (Moorthy, Choi, Bovik & De Veciana, 2012a; Moorthy, Choi, deVeciana & Bovik, 2012b; Moorthy, Choi, deVeciana & Bovik, 2012c) consists of 12 reference videos with frame rates of 30 fps and a spatial resolution of 1280×720 pixels over which different distortion types are applied to produce distorted videos. The existing distortion types are: H.264 compression at four different rates; wireless channel packet-loss distortion; freeze-frames where a loss of temporal continuity exists after freeze and where no such loss of temporal continuity takes place (not used in the analyses); rate adaptation distortion (i.e. compression rate dynamically varies between two compression rates); and temporal dynamics (compression rate is varied between several rates with different rate-switching structures). Subjective measurements acquired through viewing the different videos on a small mobile screen are available.

The LIVE-VQA dataset (Seshadrinathan, Soundararajan, Bovik & Cormack, 2010a; Seshadrinathan, Soundararajan, Bovik & Cormack, 2010b) consists of 10 reference videos with a frame rate of 25 fps or 50 fps and a spatial resolution of 768 × 432 pixels. The existing distortion types are MPEG2 compression, H.264 compression, simulated transmission through error-prone IP networks and simulated transmission through error-prone wireless networks.

Finally, the VQEG-HD3 dataset (Video Quality Experts Group, 2010) consists of 13 reference videos with a frame rate of 30 fps and a spatial resolution of 1920 × 1080 pixels. Distorted videos are achieved by applying the distortion levels hrc04, hrc07, hrc16, hrc17, hrc18, hrc19, hrc20 and hrc21.

### 3.4. Evaluation of INRF-IQA and INRF-VQA models

Existing subjective image quality and video quality scores in each of the datasets are respectively correlated with our INRF-IQA and INRF-VQA metrics. In the case of INRF-IQA, SRCC is calculated. For INRF-VQA, SRCC and PLCC are obtained. Before calculating PLCC, predicted objective video quality scores are passed through a four-parameter sigmoid function as described in Antkowiak et al. (2000):

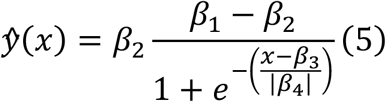

where *x* stands for the raw INRF-VQA scores, and*β*_1_, *β*_2_, *β*_3_ and *β*_4_ are its parameters. *ŷ*

## 4. RESULTS AND COMPARISONS

### 4.1. IQA

Performance of our INRF-IQA metric is evaluated on images from the LIVE (Sheikh et al., 2006), CSIQ and TID2013 (Ponomarenko et al., 2015) datasets, and compared with that of other full-reference IQA methods; results are shown in Table 1. The performance of the methods PerceptNet and LPIPS is shown in three different training scenarios: 1) training performed on the ImageNet (Deng et al., 2009) and BAPSS (Zhang et al., 2018) datasets, 2) training performed only on the BAPPS dataset, and 3) training performed on the TID2008 dataset (Ponomarenko et al., 2009). None of the IQA methods was tested on a dataset on which it had been specifically trained. GMSD needs tuning of one parameter, with its value selected so as to provide maximum performance in the three datasets considered. We can see that the performance of INRF-IQA is consistently very good across all datasets, surpassing CNN-based models such as LPIPS (which has 24.7 million parameters) (Zhang et al., 2018) and DISTS (Ding et al., 2020), as well as other extensively used classical methods like NLPD (Laparra et al., 2016). Overall, the best IQA performance is observed for PerceptNet (Hepburn et al., 2020) when its 36.3 thousand parameters are optimized for TID2008 (not for BAPPS or ImageNet), GMSD (Xue et al., 2014) and our INRF-IQA metric (with 4 parameters).

**Table 1.** Numbers indicate Spearman rank correlation coefficients (SRCC). The INRF-IQA metric is compared against a set of full-reference image quality methods: MS-SSIM (Wang et al., 2003), CW-SSIM (Wang and Simoncelli, 2005), VIF (Sheikh & Bovik, 2006), NLPD (Laparra et al., 2016), GMSD (Xue et al., 2014), MAD (Larson & Chandler, 2010), FSIM (Zhang et al., 2011), VSI (Zhang et al., 2014), DISTS (Ding et al., 2020), LPIPS (Zhang et al., 2018), and PerceptNet (Hepburn et al., 2020). LPIPS_A_ and PerceptNet_A_ are trained on the ImageNet (Deng et al., 2009) and BAPSS (Zhang et al., 2018) datasets, and LPIPS_B_ and PerceptNet_B_ are trained only on the BAPPS dataset. PerceptNetC is trained on the TID2008 dataset (Ponomarenko et al., 2009). The best three correlation values per column are marked in bold. Adapted table from Ding et al. (2021) and Hepburn et al. (2020).

|  | LIVE | CSIQ | TID2013 | (mean) |
| --- | --- | --- | --- | --- |
| MS-SSIM | 0.951 | 0.886 | 0.782 | (0.873) |
| CW-SSIM | 0.781 | 0.738 | 0.680 | (0.733) |
| VIF | **0.963** | 0.911 | 0.676 | (0.850) |
| NLPD | 0.938 | 0.937 | 0.800 | (0.892) |
| GMSD | 0.960 | **0.950** | **0.804** | **(0.905)** |
| MAD | 0.960 | **0.941** | 0.773 | (0.891) |
| FSIM | **0.963** | 0.916 | **0.802** | (0.894) |
| VSI | 0.950 | 0.923 | 0.793 | (0.889) |
| DISTS | 0.942 | 0.905 | 0.764 | (0.870) |
| LPIPS<sub>A</sub> | 0.96 | 0.93 | 0.76 | (0.883) |
| LPIPS<sub>B</sub> | 0.89 | 0.80 | 0.57 | (0.753) |
| PerceptNet<sub>A</sub> | 0.94 | 0.88 | 0.76 | (0.86) |
| PerceptNet<sub>B</sub> | 0.93 | 0.84 | 0.72 | (0.83) |
| PerceptNet<sub>C</sub> | **0.98** | **0.96** | **0.87** | **(0.936)** |
| INRF-IQA | 0.947 | 0.952 | 0.802 | **(0.900)** |

### 4.2. VQA

We start evaluating the performance of our INRF-VQA metric on videos from the very recent (and challenging) LIVE-YT-HFR dataset (Madhusudana et al., 2021). This dataset consists of reference videos of 120 fps frame rate for which distorted versions are generated by reducing their frame rate and applying different compression levels. Subjective measurements of video quality are provided for each of the videos.

It is important to note that our INRF-VQA metric, as well as many other full-reference metrics, needs reference and distorted video sequences to have the same number of frames. For this reason, when a distorted video has a lower frame rate than the reference video, either the reference video must be downsampled to match the number of frames in the distorted video or the distorted video must be upsampled to match the reference. Madhusudana et al., 2021 (see their Table 5), who test the performance of several VQA algorithms on the LIVE-YT-HFR dataset, use naive temporal upsampling by frame duplication of distorted videos. In doing so, they argue that downsampling may introduce undesired temporal artifacts in reference videos. SSIM (Wang et al., 2004), FSIM (Zhang et al., 2011) and VMAF (Li et al., 2016) are very successful, state-of-the-art, full-reference VQA metrics, and their performance comparison against our INRF-VQA metric can be seen in Table 2. Performance is shown for different frame rates in the LIVE-YT-HFR dataset: 120 fps, 60 fps and 30 fps; and in the case of 60 fps and 30 fps, both an upsampling (results taken from Madhusudana et al., 2021) and a downsampling approach are used. Upsampling is achieved by frame duplication of distorted videos and downsampling is done through frame dropping of reference videos. This way, reference and distorted videos have the same number of frames. Spearman and Pearson correlation coefficients (SRCC and PLCC) of the objective VQA metrics with subjective video quality scores are displayed.

**Table 2.** Numbers indicate SRCC and PLCC for frame rates of 120 fps, 60 fps and 30 fps in the LIVE-YT-HFR dataset. The INRF-VQA metric is compared against SSIM (Wang et al., 2004), FSIM (Zhang et al., 2011) and VMAF (Li et al., 2016). For 60 fps and 30 fps, SSIM, FSIM, VMAF and INRF-VQA results are shown both using frame duplication of distorted videos (SSIM_dup_, FSIM_dup_, VMAF_dup_ and INRF-VQA_dup_ respectively) and frame dropping of reference videos (SSIM_drop_, FSIM_drop_, VMAF_drop_ and INRF-VQA_drop_). SSIM_dup_, FSIM_dup_ and VMAF_dup_ results are taken from Madhusudana et al. (2021), their Table 5.

|  | 120 fps |  | 60 fps |  | 30 fps |  |
| --- | --- | --- | --- | --- | --- | --- |
|  | SRCC | PLCC | SRCC | PLCC | SRCC | PLCC |
| **$SSIM_{dup}$** | 0.7485 | 0.6726 | 0.2123 | 0.1845 | 0.1108 | 0.0816 |
| **$SSIM_{drop}$** |  |  | 0.5500 | 0.5077 | 0.5718 | 0.5844 |
| **$FSIM_{dup}$** | 0.7053 | 0.6368 | 0.3450 | 0.2776 | 0.2487 | 0.1786 |
| **$FSIM_{drop}$** |  |  | 0.6197 | 0.6585 | 0.5984 | 0.7524 |
| **$VMAF_{dup}$** | 0.7943 | 0.7844 | 0.5408 | 0.6015 | 0.2855 | 0.3740 |
| **$VMAF_{drop}$** |  |  | 0.6664 | 0.8122 | 0.5981 | 0.8713 |
| **$INRF-VQA_{dup}$** | 0.8507 | 0.8261 | 0.2983 | 0.2905 | 0.1198 | 0.2324 |
| **$INRF-VQA_{drop}$** |  |  | 0.7525 | 0.8262 | 0.6331 | 0.8750 |

For 60 fps and 30 fps, and for the four VQA methods, performance is seen to improve when a frame dropping strategy rather than a duplication one is used (for which correlations are low). For this reason, we use this approach as our preferred choice to evaluate INRF-VQA for 30 fps and 60 fps.

Table 3 shows a performance comparison of our INRF-VQA metric (using frame dropping when 60 fps and 30 fps videos are evaluated) against other state-of-the-art full-reference VQA metrics. SRCC and PLCC results of the objective VQA metrics with subjective video quality scores are shown for different frame rates in the LIVE-YT-HFR dataset.

**Table 3.**
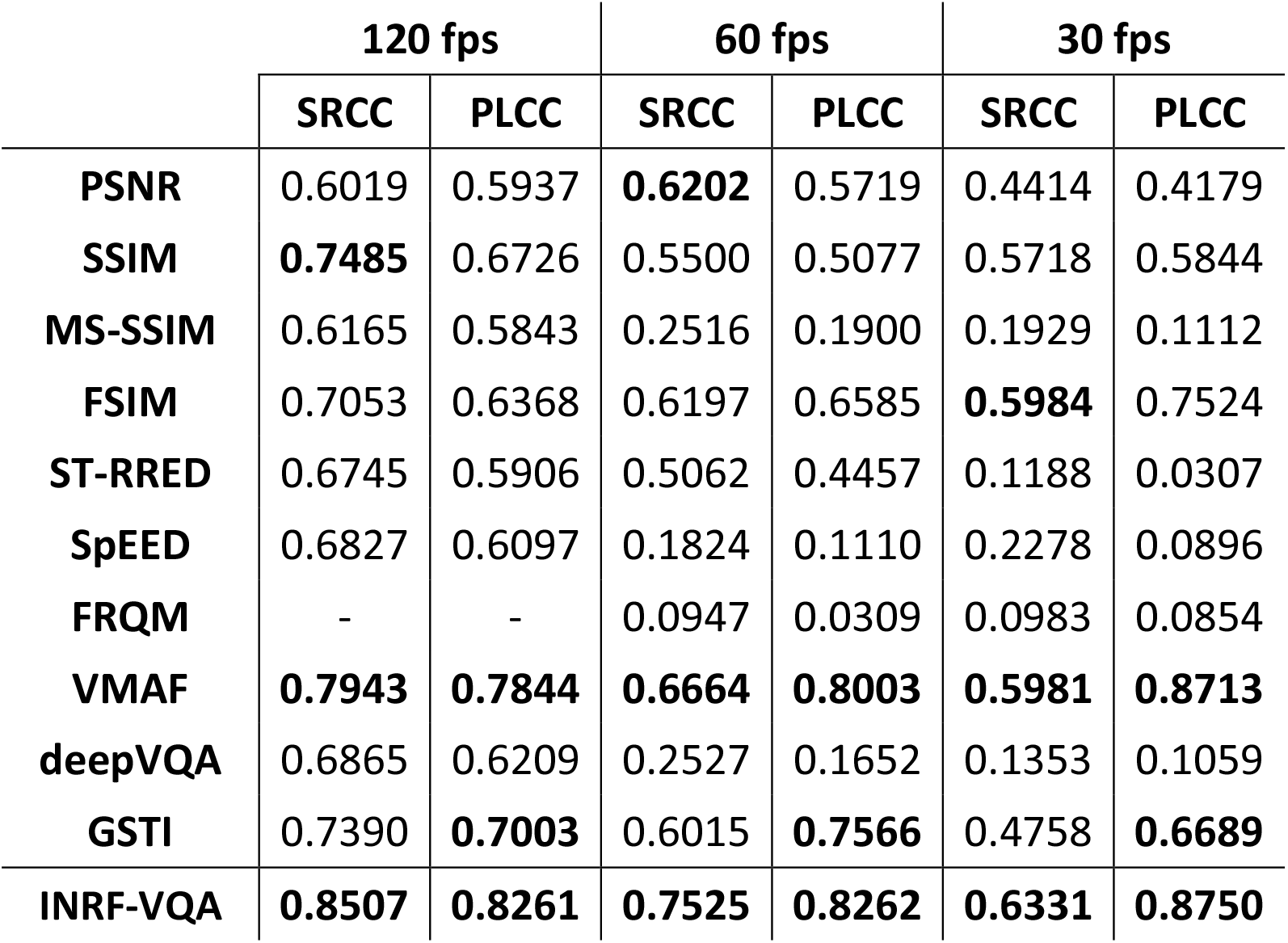
Numbers indicate SRCC and PLCC for frame rates of 120 fps, 60 fps and 30 fps in the LIVE-YT-HFR dataset. The INRF-VQA metric is compared against a set of full-reference video quality methods: PSNR, SSIM (Wang et al., 2004), MS-SSIM (Wang et al., 2003), FSIM (Zhang et al., 2011), ST-RRED (Soundararajan & Bovik, 2012), SpEED (Bampis et al., 2017), FRQM (Zhang et al., 2017), VMAF (Li et al., 2016), deepVQA (Kim et al., 2018) and GSTI (Madhusudana et al., 2020). For 60 fps and 30 fps, results for all VQA algorithms are shown using naive temporal upsampling of distorted videos except for INRF-VQA, SSIM, FSIM and VMAF, where frame dropping of reference videos is used. The best three correlation values per column are marked in bold. Adapted from Madhusudana et al. (2021).

The results in Table 3 demonstrate that INRF-VQA consistently outperforms all state-of-the-art algorithms for all frame rates tested, including the highest frame rate of 120 fps (for a fair comparison, results for SSIM, FSIM and VMAF for 60 fps and 30 fps were also computed using frame dropping, and we do not rule out that other methods could also benefit from carrying out the validation in this manner).

Next, we evaluate our INRF-VQA metric on a very popular dataset, the LIVE-MOBILE (Moorthy et al., 2012a; Moorthy et al., 2012b; Moorthy et al., 2012c). Table 4 shows, for several popular metrics and for INRF-VQA, the results for the different distortion types as well as the overall performance. As we can see, INRF-VQA ranks among the best-performing metrics for most distortions.

**Table 4.** Numbers indicate SRCC and PLCC for the different distortion types in the LIVE-MOBILE dataset: Compression, Rate Adaptation, Temporal Dynamics and Wireless; as well as global performance. The INRF-VQA metric is compared against a set of full-reference video quality methods: PSNR, SSIM (Wang et al., 2004), MS-SSIM (Wang et al., 2003), FSIM (Zhang et al., 2011), SpEED (Bampis et al., 2017), VSNR (Chandler & Hemami, 2007), VIF (Sheikh and Bovik, 2006), UQI (Wang & Bovik, 2002), NQM (Damera-Venkata et al., 2000), Weighted Signal-to-Noise Ratio (WSNR), Sinal-to-Noise Ratio (SNR), VQM (Pinson & Wolf, 2004) and MOVIE (Seshadrinathan & Bovik, 2009). The best three correlation values per column are marked in bold. Adapted from Moorthy et al. (2012), except for FSIM and SpEED values.

|  | Compression |  | Rate Adaptation |  | Temporal Dynamics |  | Wireless |  | All |  |
| --- | --- | --- | --- | --- | --- | --- | --- | --- | --- | --- |
|  | SRCC | PLCC | SRCC | PLCC | SRCC | PLCC | SRCC | PLCC | SRCC | PLCC |
| PSNR | 0.8185 | 0.7841 | 0.5981 | 0.5364 | 0.3717 | 0.4166 | 0.7925 | 0.7617 | 0.6780 | 0.6909 |
| SSIM | 0.7092 | 0.7475 | 0.6303 | 0.6120 | 0.3429 | 0.3924 | 0.7246 | 0.7307 | 0.6498 | 0.6637 |
| MS-SSIM | 0.8044 | 0.7664 | **0.7378** | **0.7089** | **0.3974** | 0.4068 | 0.8128 | 0.7706 | 0.7425 | 0.7077 |
| FSIM | **0.8896** | 0.8368 | 0.6269 | 0.5223 | 0.3020 | 0.2674 | **0.8849** | 0.8421 | 0.7491 | 0.7230 |
| SpEED | **0.9418** | **0.9379** | **0.7262** | **0.7807** | **0.4214** | **0.5697** | **0.9390** | **0.9391** | **0.8051** | **0.8118** |
| VSNR | 0.8739 | 0.8489 | 0.6735 | 0.6581 | 0.3170 | **0.4269** | 0.8559 | 0.8493 | **0.7517** | 0.7592 |
| VIF | 0.8613 | **0.8826** | 0.6388 | 0.6643 | 0.1242 | 0.1046 | 0.8739 | **0.8979** | 0.7439 | **0.7870** |
| UQI | 0.5621 | 0.5794 | 0.4299 | 0.2929 | 0.0296 | 0.2546 | 0.5756 | 0.7412 | 0.4894 | 0.6619 |
| NQM | 0.8499 | 0.8318 | 0.6775 | 0.6772 | 0.2383 | 0.3646 | **0.8985** | 0.8738 | 0.7493 | 0.7622 |
| WSNR | 0.7817 | 0.7558 | 0.5598 | 0.5365 | 0.0942 | 0.0451 | 0.7510 | 0.7276 | 0.6267 | 0.6320 |
| SNR | 0.7073 | 0.6501 | 0.5565 | 0.3988 | 0.2029 | 0.0839 | 0.6959 | 0.6052 | 0.5836 | 0.5189 |
| VQM | 0.7717 | 0.7816 | 0.6475 | 0.5910 | 0.3860 | 0.4066 | 0.7758 | 0.7909 | 0.6945 | 0.7023 |
| MOVIE | 0.7738 | 0.8103 | **0.7198** | **0.6811** | 0.1578 | 0.2436 | 0.6580 | 0.7266 | 0.6420 | 0.7157 |
| INRF-VQA | **0.8891** | **0.8868** | 0.7137 | 0.6149 | **0.5314** | **0.5834** | 0.8840 | **0.8916** | **0.7701** | **0.7684** |

**Table 5.**
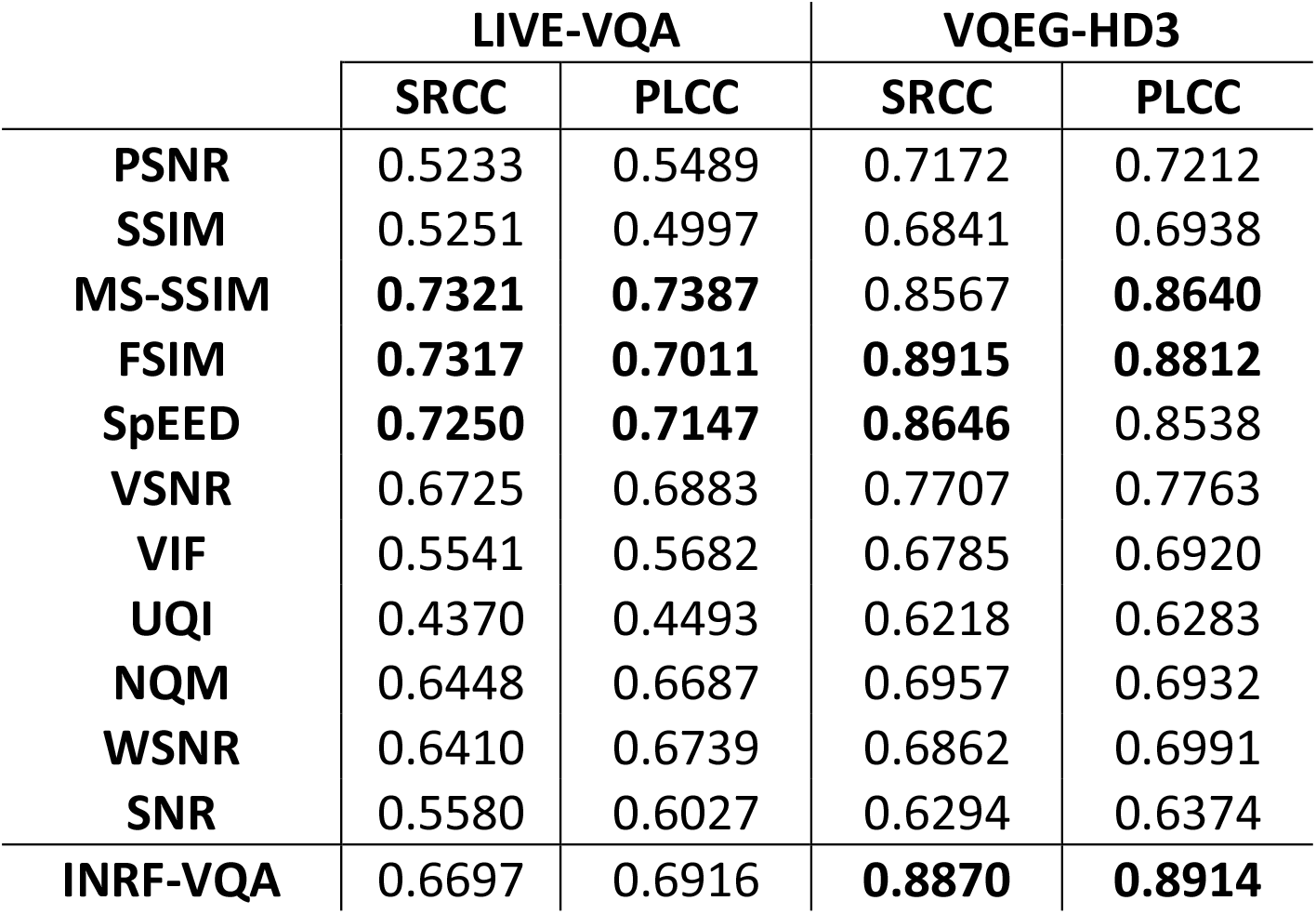
Numbers indicate SRCC and PLCC results in the LIVE-VQA (Seshadrinathan et al., 2010a; Seshadrinathan et al., 2010b) and VQEG-HD3 (Video Quality Experts Group, 2010) datasets. The INRF-VQA metric is compared against a set of full-reference video quality methods obtained from the Metrix Mux toolbox (Murthy & Karam, 2010): PSNR, SSIM (Wang et al., 2004), MS-SSIM (Wang et al., 2003), FSIM (Zhang et al., 2011), SpEED (Bampis et al., 2017), VSNR (Chandler & Hemami, 2007), VIF (Sheikh and Bovik, 2006), UQI (Wang & Bovik, 2002), NQM (Damera-Venkata et al., 2000), Weighted Signal-to-Noise Ratio (WSNR) and Signal-to-Noise Ratio (SNR). The best three correlation values per column are marked in bold.

To further test our INRF-VQA metric, we evaluate its performance on two other popular video quality datasets widely used in the VQA literature, LIVE-VQA (Seshadrinathan et al., 2010) and VQEG-HD3 (Video Quality Experts Group, 2010). Table 5 shows the correlation values in these datasets for INRF-VQA and for several popular metrics (PSNR, SSIM, MS-SSIM, VSNR, VIF, UQI, NQM, WSNR and SNR are available in the Metrix Mux toolbox [Murthy & Karam, 2010]). As we can see from these results, INRF-VQA performs very well in VQEGHD3 and shows a competitive performance in LIVE-VQA.

As a final note, we would like to remark that our INRF-VQA metric had its parameters optimized on an image dataset rather than a video dataset: this is also the approach followed by several of the best-performing VQA methods, like MS-SSIM, VIF, VSNR or NQM, while other excellent VQA algorithms are specifically trained on video quality datasets, like VMAF or GSTI.

## 5 DISCUSSION

In this study we have taken a recent neural summation model and used it as a foundation for novel metrics for image and video quality assessment. To the best of our knowledge this is a novel approach, that might pave the way for other neuroscience models to inform the creation of IQA and VQA methods.

Our validation, on popular datasets of observer scores, shows that our proposed metrics for IQA and VQA compare very well with the state-of-the-art and, very importantly, that their performance is very good and does not drop substantially for different datasets, unlike what many methods are prone to do and is often the case with those based on deep learning techniques.

For the very recent, and challenging, video quality dataset LIVE-YT-HFR (Madhusudana et al., 2021), our metric for VQA is shown to outperform all state-of-the-art models, often by a wide margin, for all frame rates considered, including a high frame rate of 120fps. Arguably, the distortions caused by the changes in frame rate in the LIVE-YT-HFR dataset are much less perceptually relevant than the artifacts created by compression: this would explain why the full-reference metrics not specifically designed to work with different frame rates (such as SSIM, FSIM and INRF-VQA) correlate well with the observers’ responses even when the information about the temporal differences is reduced (i.e. when reference frames are dropped); on the other hand, frame dropping may prevent the metrics to fully capture the potential impact of different frame rates on the perceived quality. However, we also believe that the use of frame duplication may have implications for the performance evaluation of different methods. When distorted videos have a lower framerate than reference ones, and their number of frames is matched through temporal upsampling by frame duplication, duplicated frames from the distorted video are compared to frames from the reference video that are spatially shifted one from another (whenever there is motion in a video). Methods like PSNR, with a limited performance, may indicate perceptual differences in these cases that may not be especially sensitive to the aforementioned spatial shift. However, better performing algorithms like FSIM, SSIM, VMAF or INRF-VQA itself, are sensitive towards this spatial shift. This translates into perceptual frame-difference judgements that, although accurate, are solely the result of the frame duplication strategy, and do not reflect human perception of quality. In short, the lower sensitivity of PSNR to the spatial shifts introduced by upsampling may paradoxically translate into a better performance than that of state-of-the-art algorithms which are sensitive to the artificially introduced spatial shifts.

It is important to remark that the parameters of our proposed INRF-VQA metric were optimized for image data, i.e. INRF-VQA was not trained on any video dataset. INRF-VQA is a straightforward extension of the INRF-IQA method, in which metric values are computed on a frame-by-frame basis and then averaged over time to produce a single score for the video: the fact that this simple temporal extension of an IQA method works so remarkably well for VQA challenges a common assumption in the literature, where it is thought that the best VQA metrics must be developed specifically for video (Madhusudana et al. (2021)).

Regarding future work, we want to point out that for INRF-VQA we have resorted to very simple and very effective design choices, like mean average for temporal pooling instead of a more optimal strategy (e.g. Rimac-Drlje, Vranjes & Zagar, 2009), or the fact that computations on a present frame are not influenced by past frames. For this reason, we believe that embedding temporal processing into our INRF-VQA model in a way that is more biologically realistic could prove better. As well, we would like to deepen into the study of how stacking several INRF modules produces an increase in quality prediction performance. Positive IQA results in this regard have already been reported in Bertalmío et al. (2020). We are also interested in extending both INRF-IQA and INRF-VQA so that they consider color and can be applied to high dynamic range (HDR), wide color gamut (WCG) and 8K imagery.

## 8. Author Contributions

RL and MB wrote the manuscript. RL did the final version of the INRF-VQA code and carried out the INRF-VQA experiments. ZL did the INRF-IQA version of the code and carried out the INRF-IQA experiments. MB conceptualized the original idea. All authors reviewed the manuscript.

## 10. Acknowledgments

The authors would like to thank Adrián Martín for providing us with a fast implementation of the INRF code; Jesús Malo for providing the TID2008 and TID2013 datasets; Pavan C. Madhusudana, for facilitating data from the LIVE-YouTube-HFR dataset and for his explanations about it; and Yasuko Sugito for her helpful suggestions on running VMAF.

## REFERENCES

Antkowiak, J., Baina, T. J., Baroncini, F. V., Chateau, N., FranceTelecom, F., Pessoa, A. C. F., … & Philips, F. (2000). Final report from the video quality experts group on the validation of objective models of video quality assessment march 2000.

Bampis, C. G., Gupta, P., Soundararajan, R., & Bovik, A. C. (2017). SpEED-QA: Spatial efficient entropic differencing for image and video quality. IEEE signal processing letters, 24(9), 1333–1337.

Bertalmío, M. (2019). Vision Models for High Dynamic Range and Wide Colour Gamut Imaging: Techniques and Applications (Academic Press)

Bertalmío, M., Gomez-Villa, A., Martin, A., Vazquez-Corral, J., Kane, D., and Malo, J. (2020). Evidence for the intrinsically nonlinear nature of receptive fields in vision. Scientific reports 10, 1–15. Nature Publishing Group

Betz, T., Shapley, R., Wichmann, F. A., and Maertens, M. (2015). Testing the role of luminance edges in White’s illusion with contour adaptation. Journal of vision 15, 14–14

Carandini, M., Demb, J. B., Mante, V., Tolhurst, D. J., Dan, Y., Olshausen, B. A., et al. (2005). Do we know what the early visual system does? Journal of Neuroscience 25, 10577–10597

Cavanaugh, J. R., Bair, W., and Movshon, J. A. (2002). Selectivity and spatial distribution of signals from the receptive field surround in macaque V1 neurons. Journal of neurophysiology 88, 2547–2556

Chandler, D. M., & Hemami, S. S. (2007). VSNR: A wavelet-based visual signal-to-noise ratio for natural images. IEEE transactions on image processing, 16(9), 2284–2298

Coen-Cagli, R., Dayan, P., and Schwartz, O. (2012). Cortical surround interactions and perceptual salience via natural scene statistics. PLoS computational biology 8, e1002405

Damera-Venkata, N., Kite, T. D., Geisler, W. S., Evans, B. L., & Bovik, A. C. (2000). Image quality assessment based on a degradation model. IEEE transactions on image processing, 9(4), 636–650

Deng, J., Dong, W., Socher, R., Li, L. J., Li, K., & Fei-Fei, L. (2009, June). Imagenet: A large-scale hierarchical image database. In 2009 IEEE conference on computer vision and pattern recognition (pp. 248–255).

Ding, K., Ma, K., Wang, S., and Simoncelli, E. P. (2020). Image quality assessment: Unifying structure and texture similarity. CoRR abs/2004.07728

Ding, K., Ma, K., Wang, S., and Simoncelli, E. P. (2021). Comparison of full-reference image quality models for optimization of image processing systems. International Journal of Computer Vision, 244, 1258–1281

Fukushima, Y., Hara, K., & Kimura, M. (1986). A spatio-temporal model of ganglion cell receptive field in the cat retina. Biological cybernetics, 54(2), 91–98.

Henning, G. B., Hertz, B. G., and Broadbent, D. (1975). Some experiments bearing on the hypothesis that the visual system analyses spatial patterns in independent bands of spatial frequency. Vision research 15, 247, 887–897

Hepburn, A., Laparra, V., Malo, J., McConville, R., and Santos-Rodriguez, R. (2020). Perceptnet: A human visual system inspired neural network for estimating perceptual distance. In 2020 IEEE International Conference on Image Processing (ICIP) (IEEE), 121–125

Jansen, M., Jin, J., Li, X., Lashgari, R., Kremkow, J., Bereshpolova, Y., et al. (2018). Cortical balance between on and off visual responses is modulated by the spatial properties of the visual stimulus. Cerebral Cortex 29, 336–355

Kane, D. and Bertalmío, M. (2019). A reevaluation of Whittle (1986, 1992) reveals the link between detection thresholds, discrimination thresholds, and brightness perception. Journal of vision 19, 16–16

Kaplan, E., & Benardete, E. (2001). The dynamics of primate retinal ganglion cells. Progress in brain research, 134, 17–34.

Kim, W., Kim, J., Ahn, S., Kim, J., & Lee, S. (2018). Deep video quality assessor: From spatiotemporal visual sensitivity to a convolutional neural aggregation network. In Proceedings of the European Conference on Computer Vision (ECCV) (pp. 219–234).

Kremkow, J., Jin, J., Komban, S. J., Wang, Y., Lashgari, R., Li, X., et al. (2014). Neuronal nonlinearity explains greater visual spatial resolution for darks than lights. Proceedings of the National Academy of Sciences, 201310442

Laparra, V., Ballé, J., Berardino, A., & Simoncelii, E. P. (2016). Perceptual image quality assessment using a normalized Laplacian pyramid. In Human Vision and Electronic Imaging 2016, HVEI 2016 (pp. 43–48). Society for Imaging Science and Technology.

Larson, E. C., & Chandler, D. M. (2010). Most apparent distortion: full-reference image quality assessment and the role of strategy. Journal of electronic imaging, 19(1), 011006–011006.

Li, Z., Aaron, A., Katsavounidis, I., Moorthy, A., & Manohara, M. (2016). Toward a practical perceptual video quality metric. The Netflix Tech Blog, 6(2).

London, M. and Häusser, M. (2005). Dendritic computation. Annu. Rev. Neurosci. 28, 503–532

Madhusudana, P. C., Birkbeck, N., Wang, Y., Adsumilli, B., & Bovik, A. C. (2020). Capturing video frame rate variations via entropic differencing. IEEE Signal Processing Letters, 27, 1809–1813.

Madhusudana, P. C., Yu, X., Birkbeck, N., Wang, Y., Adsumilli, B., and Bovik, A. C. (2021). Subjective and objective quality assessment of high frame rate videos. IEEE Access 9, 108069–108082

Moorthy, A. K., Choi, L. K., Bovik, A. C., & De Veciana, G. (2012a). Video quality assessment on mobile devices: Subjective, behavioral and objective studies. IEEE Journal of Selected Topics in Signal Processing, 6(6), 652–671

Moorthy, A. K., Choi, L. K., De Veciana, G., & Bovik, A. (2012b, June). Mobile video quality assessment database. In IEEE ICC Workshop on Realizing Advanced Video Optimized Wireless Networks (Vol. 6, No. 6, pp. 652–671).

Moorthy, A. K., Choi, L. K., De Veciana, G., & Bovik, A. C. (2012c, January). Subjective analysis of video quality on mobile devices. In Sixth International Workshop on Video Processing and Quality Metrics for Consumer Electronics (VPQM), Scottsdale, Arizona.

Mukherjee, D., Han, J., Bankoski, J., Bultje, R., Grange, A., Koleszar, J., Wilkins, P., & Xu, Y. (2015). A technical overview of vp9—the latest open-source video codec. SMPTE Motion Imaging Journal, 124(1), 44–54.

Murthy, A. V., & Karam, L. J. (2010, June). A MATLAB-based framework for image and video quality evaluation. In 2010 Second International Workshop on Quality of Multimedia Experience (QoMEX) (pp. 242–247). IEEE.

Peli, E. (1990). Contrast in complex images. JOSA A, 7(10), 2032–2040

Olshausen, B. A. and Field, D. J. (2005). How close are we to understanding V1? Neural computation 17, 1665–1699

Pinson, M. H., & Wolf, S. (2004). A new standardized method for objectively measuring video quality. IEEE Transactions on broadcasting, 50(3), 312–322.

Poirazi, P., Brannon, T., & Mel, B. W. (2003). Pyramidal neuron as two-layer neural network. Neuron, 37(6), 989–999.

Polsky, A., Mel, B. W., and Schiller, J. (2004). Computational subunits in thin dendrites of pyramidal cells. Nature neuroscience 7, 621

Ponomarenko, N., Jin, L., Ieremeiev, O., Lukin, V., Egiazarian, K., Astola, J., et al. (2015). Image database tid2013: Peculiarities, results and perspectives. Signal Processing: Image Communication 30, 57–77

Ponomarenko, N., Lukin, V., Zelensky, A., Egiazarian, K., Carli, M., and Battisti, F. (2009). TID2008 - A Database for Evaluation of Full-Reference Visual Quality Assessment Metrics. Advances of Modern Radioelectronics 10, 30–45

Reid, R. C., & Shapley, R. M. (2002). Space and time maps of cone photoreceptor signals in macaque lateral geniculate nucleus. Journal of Neuroscience, 22(14), 6158–6175.

Rodrigues, Y. E., Tigaret, C. M., Marie, H., O’Donnell, C., and Veltz, R. (2021). A stochastic model of hippocampal synaptic plasticity with geometrical readout of enzyme dynamics. bioRxiv

ITU-T. (2005). User Requirements for Objective Perceptual Video Quality Measurements in Digital Cable Television. ITU-T Recommendation J.143, Recommendations of the ITU, Telecommunication Standardization Sector.

Rimac-Drlje, S., Vranjes, M., & Zagar, D. (2009, May). Influence of temporal pooling method on the objective video quality evaluation. In 2009 IEEE International Symposium on Broadband Multimedia Systems and Broadcasting (pp. 1–5). IEEE.

Seshadrinathan, K., & Bovik, A. C. (2007). A structural similarity metric for video based on motion models. In 2007 IEEE International Conference on Acoustics, Speech and Signal Processing-ICASSP’07 (Vol. 1, pp. I–869). IEEE.

Seshadrinathan, K., & Bovik, A. C. (2009). Motion tuned spatio-temporal quality assessment of natural videos. IEEE transactions on image processing, 19(2), 335–350.

Seshadrinathan, K., Soundararajan, R., Bovik, A. C., & Cormack, L. K. (2010a). Study of subjective and objective quality assessment of video. IEEE transactions on Image Processing, 19(6), 1427–1441

Seshadrinathan, K., Soundararajan, R., Bovik, A. C., & Cormack, L. K. (2010b, February). A subjective study to evaluate video quality assessment algorithms. In Human Vision and Electronic Imaging XV (Vol. 7527, pp. 128–137). SPIE.

Sheikh, H. and Bovik, A. (2006). Image information and visual quality. IEEE Transactions on Image Processing 15, 430–444. doi:10.1109/TIP.2005.859378

Sheikh, H. R., Sabir, M. F., & Bovik, A. C. (2006). A statistical evaluation of recent full reference image quality assessment algorithms. IEEE Transactions on image processing, 15(11), 3440–3451.

Soundararajan, R., & Bovik, A. C. (2012). Video quality assessment by reduced reference spatiotemporal entropic differencing. IEEE Transactions on Circuits and Systems for Video Technology, 23(4), 684–694.

Silver, R. A. (2010). Neuronal arithmetic. Nature Reviews Neuroscience 11, 474–489

Theunissen, F. E., David, S. V., Singh, N. C., Hsu, A., Vinje, W. E., & Gallant, J. L. (2001). Estimating spatio-temporal receptive fields of auditory and visual neurons from their responses to natural stimuli. Network: Computation in Neural Systems, 12(3), 289.

Video Quality Experts Group. (2010). Report on the validation of video quality models for high definition video content. http://www.vqeg.org/

Vilankar, K. P. and Field, D. J. (2017). Selectivity, hyperselectivity, and the tuning of V1 neurons. Journal of vision 17, 9–9

Wang, Z., & Bovik, A. C. (2002). A universal image quality index. IEEE signal processing letters, 9(3), 81–84

Wang, Z., Bovik, A., Sheikh, H., and Simoncelli, E. (2004). Image quality assessment: from error visibility to structural similarity. IEEE Transactions on Image Processing 13, 600–612. doi:10.1109/TIP.2003.819861

Wang, Z., Simoncelli, E., and Bovik, A. (2003). Multiscale structural similarity for image quality assessment. In The Thrity-Seventh Asilomar Conference on Signals, Systems & Computers, 2003 (IEEE), vol. 2, 1398–1402

Wang, Z. and Simoncelli, E. P. (2005). Translation insensitive image similarity in complex wavelet domain. In Proceedings of IEEE International Conference on Acoustics, Speech, and Signal Processing (IEEE), vol. 2, ii–573

Wang, Z., & Li, Q. (2007). Video quality assessment using a statistical model of human visual speed perception. JOSA A, 24(12), B61–B69.

Xue, W., Zhang, L., Mou, X., and Bovik, A. C. (2014). Gradient magnitude similarity deviation: A highly efficient perceptual image quality index. IEEE Transactions on Image Processing 23, 684–695. doi:10.1109/TIP.2013.2293423

Zhang, R., Isola, P., Efros, A. A., Shechtman, E., & Wang, O. (2018). The unreasonable effectiveness of deep features as a perceptual metric. In Proceedings of the IEEE conference on computer vision and pattern recognition (pp. 586–595).

Zhang, F., Mackin, A., & Bull, D. R. (2017). A frame rate dependent video quality metric based on temporal wavelet decomposition and spatiotemporal pooling. In 2017 IEEE International Conference on Image Processing (ICIP) (pp. 300–304). IEEE.

Zhang, L., Shen, Y., and Li, H. (2014). VSI: A visual saliency-induced index for perceptual image quality assessment. IEEE Transactions on Image Processing 23, 4270–4281. doi:10.1109/TIP.2014.2346028

Zhang, L., Zhang, L., Mou, X., and Zhang, D. (2011). FSIM: A feature similarity index for image quality assessment. IEEE Transactions on Image Processing 20, 2378–2386 doi:10.1109/TIP.2011.2109730

